# Shift in pre-existing antibiotic heteroresistance explains AST change from susceptible to resistant during patient treatment

**DOI:** 10.1101/2025.11.27.690971

**Authors:** Ben Kittleson, Carter N. Abbott, Keith S. Kaye, David S. Weiss

## Abstract

The development of resistance during antibiotic therapy can lead to treatment failure, negative outcomes, and even death. While bacterial development of resistance during therapy is observed relatively frequently, it is paradoxically thought to often be mediated by rare mutations (estimated frequency of ∼1 in 10^6 or less) and is generally viewed as an unpreventable event. Here, we describe the initial classification of an isolate of Klebsiella pneumoniae as susceptible to piperacillin/tazobactam (TZP) by standard broth microdilution (BMD) testing, yet the classification of a subsequent follow-up isolate after patient treatment as resistant, seemingly demonstrating de novo evolution of TZP resistance. However, we reveal that both isolates actually exhibit heteroresistance, a phenomenon in which an isolate harbors a minor subpopulation of resistant cells, co-existing with a majority susceptible population. We demonstrate that an increase in the frequency of the resistant subpopulation in the follow-up isolate, mediated by an increase in copy number of the SHV-1 beta-lactamase, led to this isolate being classified resistant. These data highlight how a relatively minor change in the frequency of the resistant subpopulation in a heteroresistant isolate, rather than the de novo evolution of classical resistance in which all cells exhibit resistance, can underlie the classification of resistance by diagnostic tests. This emphasizes the need for more sensitive diagnostics which can detect heteroresistance, and potentially help explain why results indicating resistance are often observed rapidly after the introduction of novel antibiotics into clinical practice.

---

The development of resistance during antibiotic therapy can lead to treatment failure, negative outcomes, and even death. While bacterial development of resistance during therapy is observed relatively frequently, it is paradoxically thought to often be mediated by rare mutations (estimated frequency of ∼1 in 10^6^ or less)^1^ and is generally viewed as an unpreventable event. Here, we describe a patient from the comparator arm of the ALLIUM clinical trial with a *Klebsiella pneumoniae* complicated urinary tract infection (cUTI) that was initially classified susceptible to piperacillin/tazobactam (TZP)^2^ by standard broth microdilution (BMD) testing (MIC: 16/4 μg/mL). However, after treatment with TZP for 7 days, the test-of-cure urine culture collected 7 days after the cessation of TZP treatment tested positive, and the resulting isolate had a significantly increased TZP MIC (>256/4 μg/mL) and was classified resistant.

We investigated the cause of this resistance development and observed that the index isolate in fact exhibited heteroresistance (HR) to TZP. HR is a phenomenon in which only a subpopulation of cells (sometimes as low as 1 in 1 million) displays increased resistance to a given antibiotic compared to the rest of the cells in the population. Due to the low frequency of resistant cells, and as in the case described here, HR is often undetected by AST tests including BMD. Further, HR has been linked to antibiotic treatment failure in patients.^3^

Population analysis profile (PAP) is the current gold standard test to detect HR and interestingly revealed that the frequency of the resistant subpopulation increased from ∼1 in 50,000 in the index isolate to ∼1 in 1,000 in the follow-up isolate (**Figure 1; S1**). Further, whole genome sequencing revealed no mutations known to be associated with beta-lactam resistance but instead detected an increase in the copy number of the SHV-1 beta-lactamase in the follow-up isolate (**Figure S2**). An increase in copy number and resulting increased production of beta-lactamases has been shown capable of mediating beta-lactam HR.^5^

**Figure 1.**
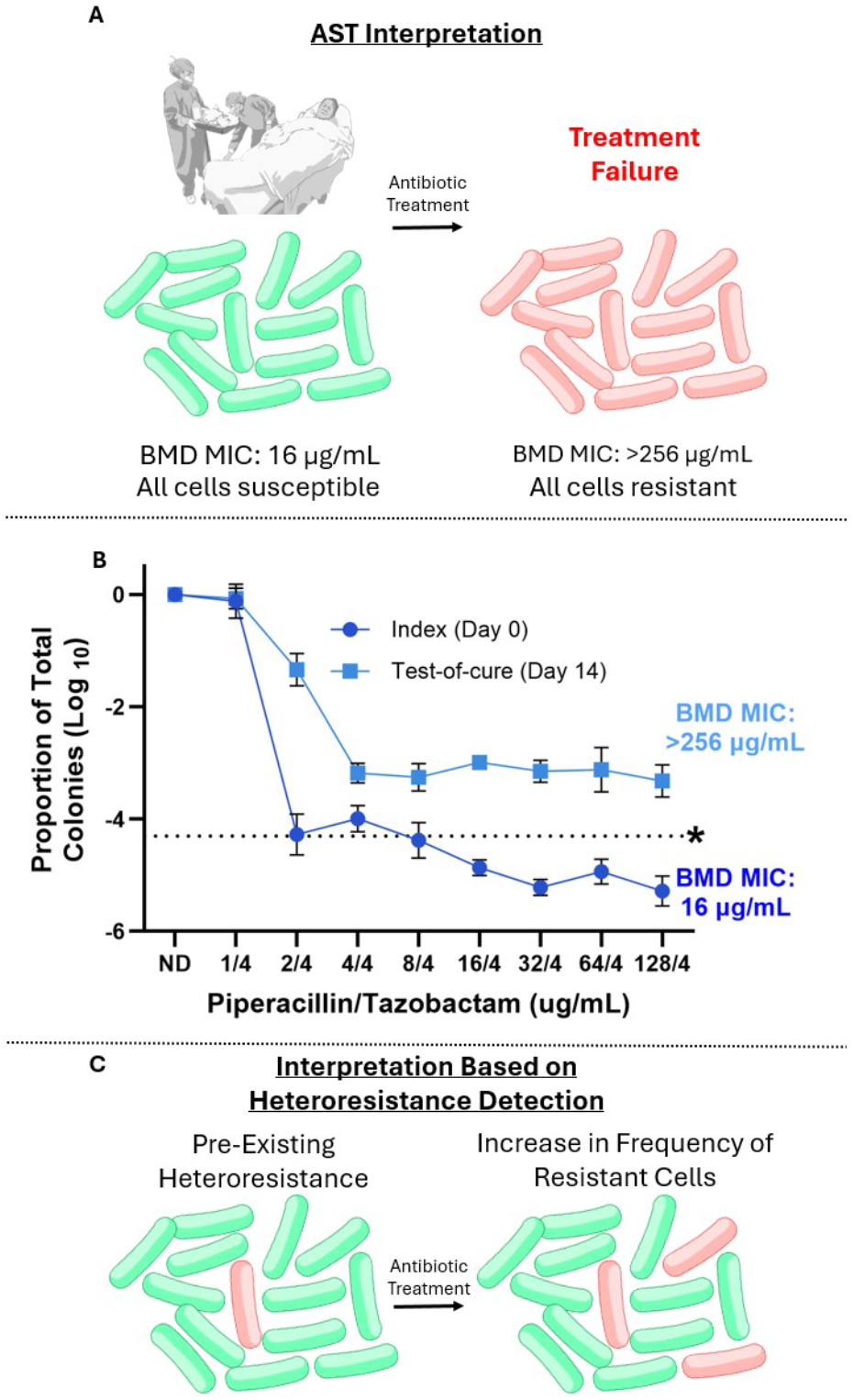
*Klebsiella pneumoniae* clinical isolate heteroresistance profile. **A)** Interpretation of the observed emergence of TZP resistance based on the clinical BMD data. **B)** Population analysis profile (PAP) results are shown for both the original and follow-up isolate. Isolates with subpopulations greater than 1*10^-6^ (1 in 1 million) cells surviving at the breakpoint concentration (32/4 μg/mL for piperacillin/tazobactam), excluding resistant isolates, were classified as heteroresistant. The dashed line represents ∼1/20,000 cells being resistant, which appears to be the approximate limit of detection for an isolate to appear resistant by BMD based on these findings, as well as previous findings related to cefiderocol.^4^ **C)** New proposed interpretation of the observed emergence of TZP resistance during treatment being the result of pre-existing heterogeneity with an elevated frequency of resistance cells after treatment.

A hallmark of HR is that the frequency of the resistant subpopulation is unstable, increasing in the presence of a relevant antibiotic yet subsequently reverting upon culture in the absence of antibiotic. Indeed, both the index and test-of-cure isolates exhibited an increase in the frequency of the resistant cells when the strains were cultured in TZP and subsequently decreased in drug-free media *in vitro*, further confirming the isolates as HR (**Figure S3**). Furthermore, the copy number of the SHV-1 beta-lactamase increased when each isolate was cultured in TZP (**Figure S2)**.

These data indicate that rather than an overwhelming, stable shift of all cells from a susceptible to a resistant phenotype, a relatively small increase in the frequency of a subpopulation of resistant cells can explain what would typically be interpreted as the novel evolution of resistance during patient therapy. This increase in the frequency of the resistant subpopulation can be mediated by an increase in the copy number of a resistance gene, demonstrating the transient flexibility that bacteria can have in responding to antibiotic stress, in the absence of *de novo*, fixed mutation. As there is currently no AST that reliably detects HR, clinicians do not have information on HR or resistant subpopulation frequency when making prescribing decisions. These results highlight that more sensitive diagnostics are an urgent need and potentially help to explain why events interpreted as novel evolution of resistance are commonly observed in the clinic.

## Notes

### Competing Interest Statement

DSW is a holder of patents related to heteroresistance.

### Funding

- ARPA-H: AY1AX000002
- NIAID: AI158080
- Veteran's Administration: BX005985
- Burroughs Wellcome Fund

