## Supplemental Figures 1-3 and Table 1 for "Shift in pre-existing antibiotic heteroresistance explains AST change from susceptible to resistant during patient treatment"

**Supplementary Appendix:**

**Table of Contents:**


| | |
|---|---|
| Supplementary Methods | 2-3 |
| Supplementary Figure 1 | 4 |
| Supplementary Figure 2 | 5 |
| Supplementary Figure 3 | 6 |
| Supplementary Table 1 | 7 |

### Supplementary Methods

Bacterial Isolates and Clinical Presentation

Clinical isolates were obtained from the randomized double-blind ALLIUM clinical trial (NCT03687255). Both the day 0 and test-of-cure isolates were obtained from a diabetic female patient under the age of 65 with no prior antibiotic therapy. The patient presented with dysuria, fever, nausea, costochondritis, flank pain, and leukopenia. Piperacillin, 4 g/tazobactam, 0.5 g was administered by a 2-hour infusion every 8 hours for 7 days. Blood cultures were negative upon admission. The patient's symptoms showed improvement after 3 days of antibiotic treatment.

Population Analysis Profile (PAP)

Population analysis profiles (PAP) were conducted as previously described[1,2]. Five to six biological replicates of each bacterial isolate were each inoculated into 1.5 mL of Mueller-Hinton broth (MHB) and incubated at 37°C overnight with shaking at 220 RPM. A 1:10 serial dilution series was then made in PBS for each replicate. 7.5 µL of each sample in the dilution series was then plated onto Mueller-Hinton agar plates containing increasing concentrations of piperacillin/tazobactam (TZP) from 1/4 µg/mL to 128/4 µg/mL, along with a no drug control. Plates were then incubated overnight at 37°C before colonies were counted and compared to the no drug control plate. Isolates were classified as heteroresistant if they exhibited between 50% and 0.0001% survival at concentrations of both 32/4 µg/mL and 64/4 µg/mL (1x and 2x the 2025 CLSI breakpoint concentration for TZP). Isolates with 50% survival or greater at the breakpoint were classified as resistant, while isolates with less than 0.0001% survival at the breakpoint concentration were classified as susceptible.

Broth Microdilution (BMD)

Three colonies were taken from an MHA plate and inoculated into MHB and incubated for 4-6 hours at 37°C with shaking at 220 RPM. Cultures were then diluted in MHB to a concentration of $10^6$ CFU/mL. 50 µL of culture was then added to a 96-well plate containing 2-fold serial dilution from 256 µg/mL to 2 µg/mL piperacillin plus 4 µg/mL tazobactam for a final concentration of 5 x $10^4$ CFU/well, along with no drug controls. Plates were then incubated statically for 16 hours at 37°C. MICs were determined via visual inspection to identify the lowest concentration of TZP without any growth. All isolates were tested with three biological replicates.

Heteroresistance Stability Assay

Cultures of bacterial isolates were prepared in 3 mL of MHB and incubated at 37°C overnight with shaking at 220 RPM. A PAP was then conducted on each isolate, before passaging the overnight cultures into 3mL of fresh MHB containing TZP (16/4 µg/mL) and incubated at 37°C overnight with shaking at 220 RPM. This process of conducting a PAP and passaging without piperacillin/tazobactam was then repeated daily for multiple days until a decreasing frequency of resistant cells at the breakpoint concentration was observed. All isolates were tested with five to six biological replicates.

Whole Genome Sequencing

Cultures were grown in 1.5 mL of MHB with a minimum of four biological replicates for samples in both the presence and absence of drug (16/4 µg/mL TZP). Bacterial genomic DNA was extracted from overnight cultures using a Wizard® Genomic DNA Purification Kit (Promega, Madison, WI). Genomes and copy number variation analyses were conducted using Illumina 400 Mbp reads and assembled against a reference genome of isolate 352 assembled via 600 Mbp Nanopore sequencing. All sequencing was performed by SeqCenter LLC (Pittsburgh, PA).

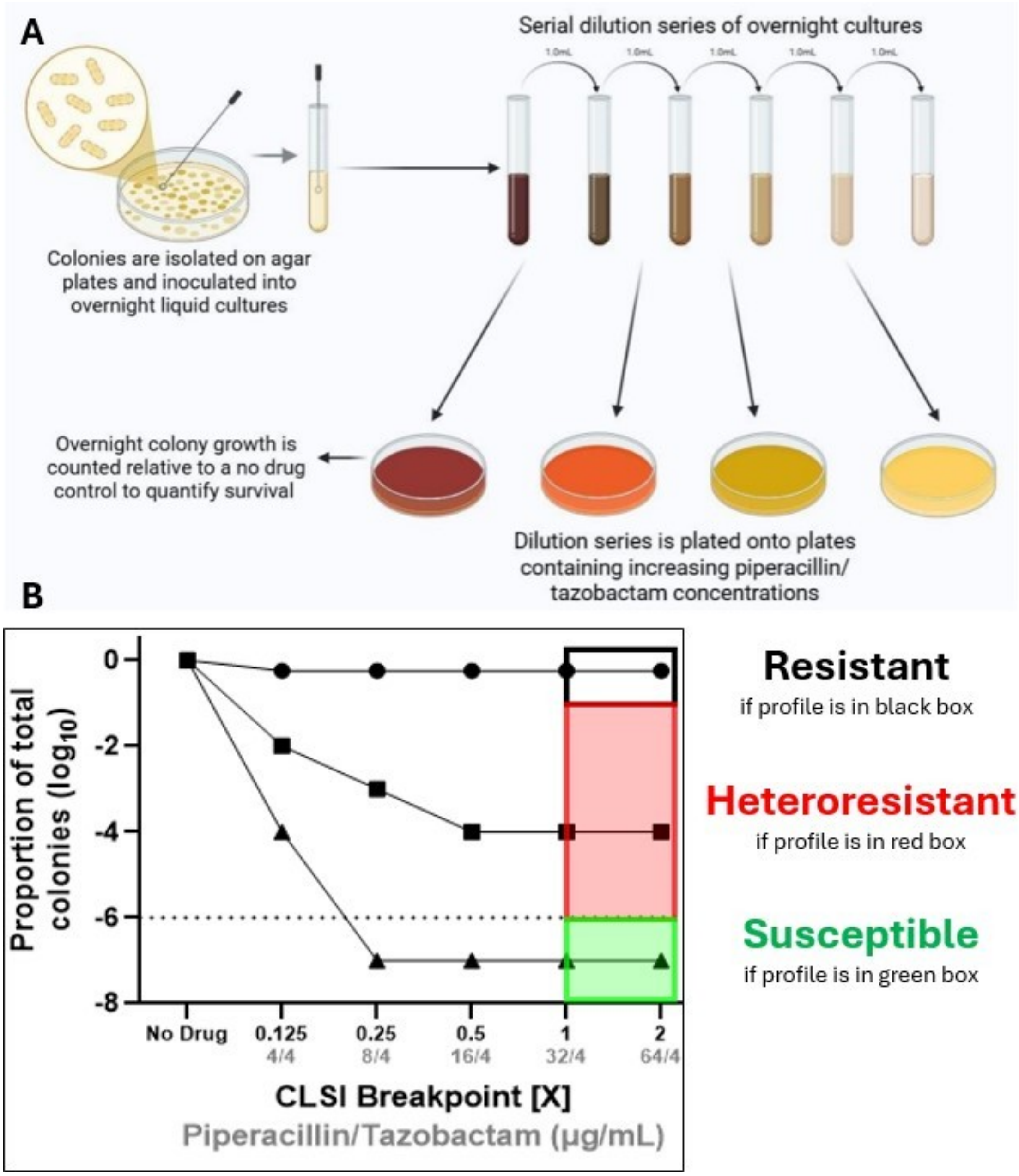


**Supplemental Figure 1. Population Analysis Profile**. **A)** Explanation of population analysis profile (PAP) methodology and **B)** categorization of isolates based on PAP results. Figure made with BioRender.

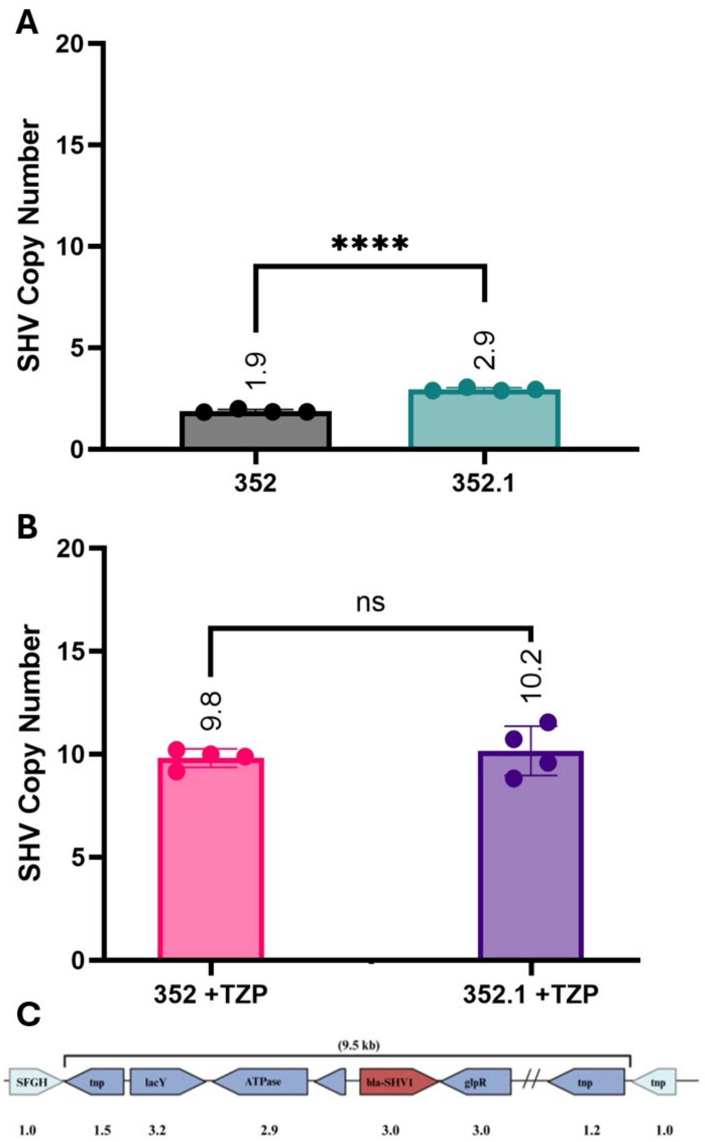


**Supplemental Figure 2. Amplification of SHV-1 β-lactamase following patient treatment with TZP.** SHV β-lactamase copy number was measured after culturing in either **A)** Mueller-Hinton broth or **B)** Mueller-Hinton broth with TZP. **C**) An abbreviated version of the amplicon containing the SHV β-lactamase, which is present in both the chromosome and the plasmid, is depicted. The numbers below each gene represent the relative copy number in the test-of-cure isolate (352.1) grown without drug.

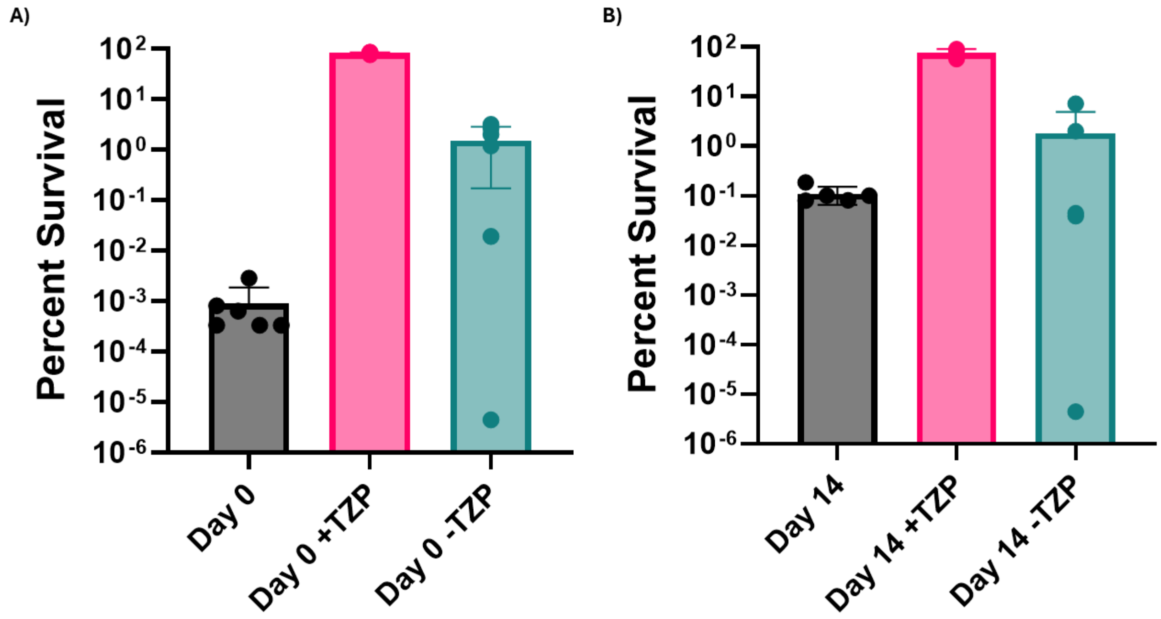


**Supplemental Figure 3. Instability of the frequency of the resistant subpopulation characterisitic of heteroresistance**. The geometric mean of percent survival at the breakpoint concentration (32/4 μg/mL TZP) is shown for the Day 0 (**A**) and Day 14 isolates (**B**) after passaging in media with 16/4 μg/mL TZP (+TZP), as well as after subsequent passaging in antibiotic-free media (-TZP) based on population analysis profile results.
